# Microbial invasions influence plant-microbiome feedback and coexistence

**DOI:** 10.1101/2025.01.31.636005

**Authors:** William H. Parker, Chase J. Rakowski, Senay Yitbarek

## Abstract

Plant-soil feedback (PSF) theory leverages the increasingly recognized ecological significance of microbiomes to provide insight into plant coexistence. However, PSF theory generally assumes that microbiomes have static composition and properties. In reality, plant communities are thought to commonly experience microbial colonization. Here, we use a PSF model to study microbial invasions of a community of two plant species and their associated microbiomes. We manipulate initial plant and microbiome evenness, microbial colonization timing, and properties of all actors, then record resulting community membership. Invasion success is higher during transient dynamics than when approaching equilibrium. Furthermore, stability analysis of a set of communities shows that invasion success is determined by the effects of the plants and the invader on each other’s growth rates. When successful, microbial invaders commonly disrupt plant coexistence. Our study demonstrates a promising framework for understanding biodiversity dynamics via integration of host-microbiome dynamics, dispersal, and time-dependent interactions.

## Introduction

Theoretical and empirical studies on feedback between host plants and their microbiomes, known as “plant-soil feedback,” highlight that microbes are important drivers of plant community structure (Bever *et al*. 2015; Mangan *et al*. 2010). Mutualists and pathogens can generate negative feedback (e.g., coexistence) or positive feedback (e.g., monodominance) during plant-plant interactions (Jiang *et al*. 2020). The ecological conditions that give rise to either positive or negative feedback in plant communities depend on the life-history traits of both microbes and plants (Holt & Lawton 2003; Mordecai 2013b, a; Parker & Gilbert 2004; Spear *et al*. 2015). However, many microbes have high dispersal rates, meaning colonization by new microbes is frequent (Chaudhary *et al*. 2022; Finlay & Clarke 1999). Additionally, variation in microbial arrival time can be an important driver of plant community structure. For example, Peay (2018) showed that delayed arrival of an ectomycorrhizal fungal partner resulted in reduced growth of its conspecific host, while benefiting its heterospecific plant competitor. This suggests that variation in microbial properties and colonization can have important implications for plant coexistence. Yet, plant-soil feedback studies have assumed microbiomes to be static in their properties and composition, and generally have only implicitly considered the temporal context of microbial responses to plant hosts (i.e. conditioning phase) and plant responses to the microbes (i.e. feedback phase; e.g., Hannula *et al*. 2020).

Over the last few decades, theoreticians have developed an influential body of work explaining the potential facilitation or constraints to plant coexistence induced by plant-soil feedback. Bever et *al.* (1997) developed a foundational model which departs from the Lotka-Volterra competition approach, instead relying on feedback between plant species and their associated microbiomes alone to determine plant coexistence outcomes. This two-plant model finds that if microbes inhibit the growth of their associated plant (i.e., negative feedback), plant coexistence is enhanced, while positive plant-soil feedback inhibits coexistence. The generalizability of this model was called into question when Miller *et al*. (2022) showed that extending the model to incorporate additional plant species almost always leads to a breakdown in coexistence. However, Abbott *et al*. (2021) allowed more robust coexistence and enhanced predictability by combining the model from Bever *et al*. (1997) with the Lotka-Volterra competition approach outlined in Bever (2003). Subsequent studies have added further realism, for example incorporating priority effects – i.e., differential outcomes based on plant species arrival (Senthilnathan & D’Andrea 2024) or germination order (Ke & Levine 2021), including for a third non-native plant species (Zou *et al*. 2024). However, the effects of microbial colonizations and their timing on plant coexistence have not yet been explored in this plant-soil feedback context.

Resident communities, including microbiomes, can prevent or facilitate the establishment of new microbes in ecosystems. Biotic interactions can be a key determinant of whether a microbe successfully colonizes a new community (Albright *et al*. 2020). Relatedly, the “biotic resistance hypothesis” in invasion ecology posits that more diverse ecosystems are more resistant to invaders as fewer niches become available, while communities with lower biodiversity are more prone to invasion (Elton, 2020; Jeschke *et al*., 2014). In microbial communities negative associations have been found between microbial biodiversity and the likelihood of new colonizations (Dooley *et al*. 2024; Eisenhauer *et al*. 2016; van Elsas *et al*. 2012), although there is also evidence for the opposite pattern in less diverse microbial communities (Madi *et al*. 2020). While the mechanisms underlying colonization resistance are not well resolved, the order and timing of colonization also play an important role in the assembly of microbial communities and ecological success of invading microbes (Debray *et al*. 2022; Litvak & Bäumler 2019). Specifically, communities undergoing transient dynamics can be more prone to colonization by new species (Dooley *et al*. 2024; Fukami & Nakajima 2011; Tucker & Fukami 2014). However, the relative importance of biotic interactions, resident diversity, and colonization timing in determining the establishment of new microbes in plant-microbiome communities remains unclear.

Here, we extend a plant-soil feedback model developed by Abbott *et al*., (2021) to explore how the invasion of a microbe can influence coexistence in a community of two host plants and their associated microbiomes. We conduct invasion simulations across a large swathe of parameter space representing colonizations of many resident communities by many different microbial invaders, and estimate the distribution of resulting final community membership. Furthermore, we explore the relationship between invasion success and initial plant and microbiome evenness as well as invasion timing. Finally, we examine the relationship between final community state and the plant-invader relationship. We find that microbial invader success is determined by the plant-invader relationship rather than microbiome or host plant evenness, and is highest when colonization occurs during transient dynamics well before the resident community reaches an equilibrium. In many of the simulated communities, invader establishment disrupts the feedback dynamics that maintain plant coexistence, though continued coexistence is also common. This study expands plant-soil feedback theory by providing an initial exploration of the conditions allowing, and consequences of, microbial invasions of plant communities.

## Material and methods

### Model

We extend a host-microbiome model developed by Abbot et al. (2021) by incorporating a non-host-associated microbial invader into the community of two host-microbiome pairs. The model developed by Abbot et al. (2021) incorporates the Lotka-Volterra competition modeling framework (Bever 2003) into the plant-soil feedback model first proposed by (Bever *et al*. 1997). The community consists of two plant hosts, *N*_1_ and *N*_2_, and their corresponding host-associated microbiomes *M*_1_ and *M*_2_, respectively. The density of each plant host positively affects the growth of its own microbiome. As in previous models (Abbott *et al*. 2021; Bever *et al*. 1997), relative proportions of the microbiomes are modeled rather than explicit microbiome densities, retaining the focus of the model on the plant populations and feedback between microbiomes rather than the size of microbial populations.

Parameter *v* represents the strength of support for *M*_2_ by its host *N*_2_ relative to the support for *M*_1_ by *N*_1_ which is assumed to have magnitude 1 (Fig. 1). The relative proportion of microbiomes feeds back on the corresponding host densities: *M*_1_ influences *N*_1_ via the parameter *α*_11_ and *M*_2_ influences *N*_2_ via *α*_22_. Additionally, the microbiomes influence the growth of their non-host plant: *M*_1_ impacts the density of plant host *N*_2_ via the parameter *α*_21_, and *M*_2_ impacts *N*_1_ via *α*_12_. The two plants *N*_1_ and *N*_2_ affect each other’s growth via interspecific competition represented by *c*_21_ and *c*_12_, and through intraspecific density dependent regulation represented by *c*_11_ and *c*_22_ (Abbott *et al*. 2021; Bever 2003; Bever *et al*. 1997). Departing from previous models, we include a microbial invader *M*_3_. *M*_3_ directly affects the densities of plant hosts *N*_1_ and *N*_2_ via the parameters *α*_13_ and *α*_23_, respectively, and *N*_1_ and *N*_2_ affect the density of *M*_3_ via the parameters *e*_1_ and *e*_2_, respectively (Fig. 1). Note that *M*_3_ is not a plant-associated microbiome, but it influences *N*_1_ and *N*_2_ and is affected by them.

**Figure 1:**
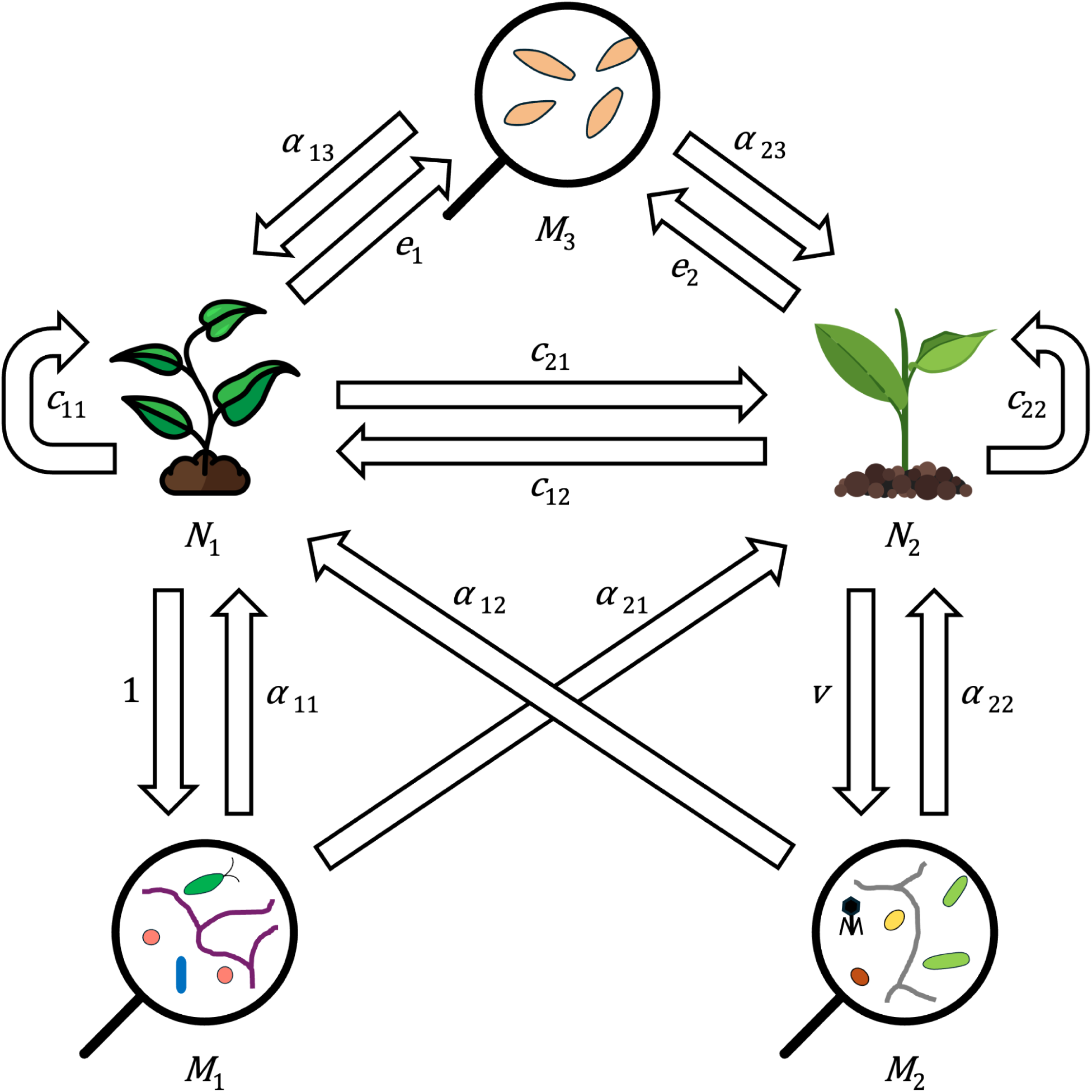
Diagram representing the state variables in the model as well as the direct interactions between them. Arrows are labeled with the key parameters that determine the strength and sign of the interaction. State variables, clockwise from bottom left: *M*_1_, the microbiome associated with host *N*_1_; *N*_1_, host 1; *M*_3_, the invading microbe; *N*_2_, host 2; and *M*_2_, the microbiome associated with host *N*_2_. *M*_1_, *M*_2_, and *M*_3_ each influence the density of both hosts, *N*_1_ and *N*_2_. Meanwhile, *N*_1_ and *N*_2_ both influence the density of only their associated microbiome and of *M*_3_ (e.g. *N*_1_ directly affects *M*_1_ and *M*_3_ but not *M*_2_). Lastly, both *N*_1_ and *N*_2_ influence the density of each other and of themselves via interspecific and intraspecific competition, respectively.

The changes in host densities and microbiome frequencies are described using components of the Lotka-Volterra competition model. The per-capita growth rate of plant host *N*_i_ is represented by the parameter *r*_i_. Interspecific and intraspecific coefficients, *c*_i_, describe density-dependent resource competition between and within plant hosts. Plant hosts *N*_1_ and *N*_2_ increase the proportion of their respective microbiomes, *M*_1_ and *M*_2_, at a rate proportional to the host’s frequency in the community. Host densities also cause changes in the frequency of the non-host associated microbe *M*_3_. Microbiome growth is a function of the relative densities of plant hosts and the product of microbiome frequencies. The strength of plant support for the frequency of *M*_3_ is quantified by *e*_i_ where positive values indicate that *N*_i_ supports *M*_3_ and negative values indicate that *N*_i_ decreases *M*_3_ relative frequency. Furthermore, *M*_3_ modifies the growth rate of both hosts. The strength of growth rate modification by *M*_3_ on *N*_i_ is quantified by the model parameter *α*_i3_, which can be positive or negative. We define the frequency of *M*_3_ implicitly in the model: *M*_3_ = 1 - (*M*_1_ + *M*_2_).

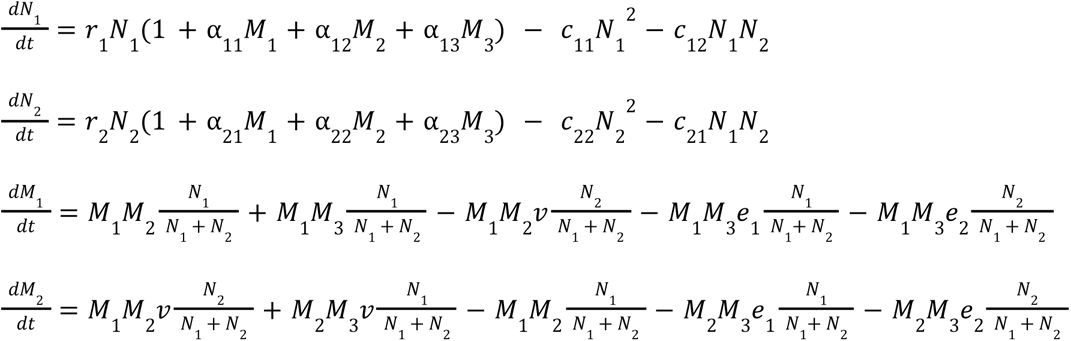

### Model Assumptions

We maintain several assumptions made by the original plant-soil feedback model proposed by Bever et al. (1997). First, we assume that microbes (i.e. *M*_1_, *M*_2_, and *M*_3_) modify plant growth proportionally to the relative frequencies of the microbes. For example, if a plant were exposed to a microbial community that is 30% *M*_1_, 40% *M*_2_, and 30% *M*_3_, its growth rate would be multiplied by a factor of 0.3*α*_i1_ + 0.4*α*_i2_ + 0.3*α*_i3_, where *α*_ij_ is the multiplicative growth rate modification of *M*_j_ on *N*_i_ when grown in a microbial community that is 100% *M*_j_. Similarly, the plants modify microbial growth rates proportionally to the plants’ relative frequencies.

Additionally, each plant host only modifies the growth of its resident microbiome and the invader – e.g., the effect of *N*_1_ on *M*_2_ is indirect via *N*_1_’s effects on *M*_1_ and *M*_3_. The microbial entities also indirectly interact via their relationships with the plants rather than interacting directly. Finally, the effects of the plants on their own microbiomes are assumed to be positive.

The assumption that microbes alter plant growth rates proportionally to their relative abundance makes it possible to model only microbial relative abundances rather than densities. This allows a reduction in the number of state variables, facilitating more in-depth analysis of the model. Microbial relative abundances are generally more applicable to empirical data, since technological constraints mean that most microbial datasets represent relative abundances more accurately than absolute abundances. Additionally, relative abundances correspond well with coexistence theory and experiments, since relative abundance is used as a metric for the stabilizing mechanism in modern coexistence theory (Adler *et al*. 2007).

### Invasion simulations

To broadly analyze the model, we first performed numerical simulations to investigate the effects of parameters, initial plant and microbiome evenness, and invasion timing on invasion outcomes. We began by using a Latin hypercube sampling method to efficiently and uniformly sample 1,000,000 parameter sets from the 11-dimensional parameter space within the ranges given in Table 1. For each of the sampled parameter sets, we used the one-plant equilibrium values for the system to calculate the equilibrium density of each plant species if it were alone. Then we set each plant at 50% of this individual equilibrium density and set the microbiome composition (*M*_1_ and *M*_2_) to be proportional to the relative frequency of the plants. From these initial conditions the simulation was run for 500 time-steps to allow the system to reach equilibrium, and the resulting densities were recorded to determine the outcomes associated with each parameter set. Those parameter sets that give rise to internal equilibria (all state variables above zero) were then used for invasion simulations.

**Table 1:** Parameters sampled in the analyses and the ranges across which they were sampled.

| Parameter | Interpretation | Minimum value<br>sampled | Maximum value<br>sampled |
| --- | --- | --- | --- |
| $r_1$ | Growth rate of $N_1$ | 0.5 | 2 |
| $r_2$ | Growth rate of $N_2$ | 0.5 | 2 |
| $\alpha_{11}$ | Frequency-dependent growth rate<br>modification of $N_1$ by $M_1$ | -2 | 2 |
| $\alpha_{12}$ | Frequency-dependent growth rate<br>modification of $N_1$ by $M_2$ | -2 | 2 |
| $\alpha_{13}$ | Frequency-dependent growth rate<br>modification of $N_1$ by $M_3$ | -2 | 2 |
| $\alpha_{21}$ | Frequency-dependent growth rate<br>modification of $N_2$ by $M_1$ | -2 | 2 |
| $\alpha_{22}$ | Frequency-dependent growth rate<br>modification of $N_2$ by $M_2$ | -2 | 2 |
| $\alpha_{23}$ | Frequency-dependent growth rate<br>modification of $N_2$ by $M_3$ | -2 | 2 |
| $c_{11}$ | Density-dependent effect of $N_1$ on $N_1$ | 0.1 | 0.5 |
| $c_{12}$ | Density-dependent effect of $N_2$ on $N_1$ | 0.1 | 0.5 |
| $c_{21}$ | Density-dependent effect of $N_1$ on $N_2$ | 0.1 | 0.5 |
| $c_{22}$ | Density-dependent effect of $N_2$ on $N_2$ | 0.1 | 0.5 |
| $v$ | Strength of support for $M_2$ by $N_2$ as a factor of the support for $M_1$ by $N_1$ | 0.1 | 10 |
| $e_1$ | Strength of support for $M_3$ by $N_1$ as a factor of the support for $M_1$ by $N_1$ | -2 | 2 |
| $e_2$ | Strength of support for $M_3$ by $N_2$ as a factor of the support for $M_1$ by $N_1$ | -2 | 2 |

In initial invasion simulations, 1000 of the parameter combinations that gave rise to internal equilibria were simulated for an additional 100 time-steps from their achieved equilibrium point with 1% of the total microbiome replaced with the invader *M*_3_. This process was repeated 1000 times per set of parameters with each invader having different characteristics sampled using the same Latin hypercube sampling method used to sample resident system characteristics. The final state of the system following these simulations was recorded and analyzed to determine the probability of any plant extinction (defined as *N*_1_ or *N*_2_ density below the 10^-4^ extinction threshold) and probability of invader persistence (*M*_3_ frequency greater than 10^-4^ extinction threshold) as a function of pre-invasion microbiome evenness (% *M*_1_) and plant evenness (% *N*_1_).

To investigate the effects of invasion timing on invasion dynamics, 1000 parameter combinations that both reached an internal equilibrium and were locally stable were selected and simulated one time-step at a time until they reached equilibrium (defined as less than a 10^-4^ unit change in all state variables over 1 time-step), and this time to equilibrium was recorded. Then the invader was introduced to these systems at the beginning of the simulation and the system was simulated until it reached its new post-invasion equilibrium. This process was repeated with the invader instead being introduced at the time-point halfway to the time point at which the system had reached equilibrium without the invader, and again at the time-point at which the system reached equilibrium. Finally, all of these steps were repeated 100 times for each parameter combination with various invader parameters sampled using the Latin hypercube sampling method.

All simulations were conducted in R version 4.1.3 (R Core Team 2022), with the *lhs* package for parameter sampling (Carnell 2022), the *deSolve* package for numerical integration (Soetaert *et al*. 2010), and the *tidyverse* (Wickham *et al*. 2019), *patchwork* (Pedersen 2025), and *ggrepel* packages for visualization (Slowikowski 2024). The *lsoda* solver was selected for its speed and dynamic switching between stiff and non-stiff methods as needed (Hindmarsh & Petzold 2005). As a further protection against numerical instability, an extinction threshold of 10^-4^ was used for all elements of the system under which any entity that drops below 10^-4^ is considered extinct and considered to have density 0 in future iterations of the simulation.

### Equilibrium invasion analysis

For the second phase of our analysis, we investigated the effects of invader characteristics on invasion outcomes by performing a targeted equilibrium invasion analysis for a focal pre-invasion community. We were not able to perform the analysis on all possible communities (i.e., parameter combinations) due to the computational complexity of the analysis and the large number of equilibria. The first step was to identify all possible equilibria of the model. To do this, we used Mathematica version 13.1 to solve for the plant host densities and microbe frequencies for which the derivatives of all state variables with respect to time are uniformly zero (Wolfram Research, Inc. 2022). We excluded equilibria with negative values as they are not biologically feasible. Then we used the Routh-Hurwitz stability criterion to determine the stability of the equilibria. This method approximates the system-governing differential equations as a linear transformation around a given equilibrium point (the Jacobian matrix of the system evaluated at that equilibrium point), then uses the signs of the eigenvalues to determine if a small perturbation from equilibrium will grow or shrink.

Next, we chose one set of resident system parameters for which the internal equilibrium is stable, {*r*_1_=0.71, *r*_2_=1, *α*_11_=1, *α*_12_=1.8, *α*_21_=0.66, *α*_22_=1, *c*_11_=1, *c*_12_=0.86, *c*_21_=1, *c*_22_=1, *v*=0.75}. This set of parameters was obtained by using Mathematica’s built-in *Solve* function to select a set of parameters for which the Routh-Hurwitz stability conditions for the two-host two-microbiome (pre-invasion) were satisfied. Then the stability of each equilibrium point was computed for given values of *α*_13_, *α*_23_ (i.e., the effects of the invader on plant growth rates), *e*_1_, and *e*_2_ (i.e. the effects of the plants on invader growth rate) to determine the long term behavior of the system given those plant-invader properties if an invader was introduced in a small dose. If the internal equilibrium remained stable, the system resisted invasion. If the internal equilibrium became unstable, the stability of other equilibria was used to determine the long-term behavior of the system. If the only stable equilibria for a given set of invader characteristics were those in which only *N*_1_ remains, this region in parameter space was recorded and the same was done for *N*_2_. These investigations were conducted for invader characteristics for every combination of *α*_13_, *α*_23_ ∈ {-1,1} and *e*_1_, *e*_2_ ∈ [-2,2]. We chose *e*_1_ and *e*_2_ to be the continuous variables of the analysis because they more directly determine whether the invader is able to persist in the system.

## Results

The microbial invader *M*_3_ successfully invades the resident community under 52% of parameter combinations tested overall. Post-invasion, the system reaches one of 15 new equilibrium communities, comprising most possible combinations of *M*_3_ and the resident entities (Fig. 2, Fig. S1). While the invader simply joins the resident community with no extinctions in 10% of successful invasions, in the remaining 90% one or more resident entities are driven extinct. Often, only one host and *M*_3_ are left in the community (54% of successful invasions). After 29% of successful invasions, both hosts remain but *M*_3_ replaces one or both resident microbiomes. Overall, the hosts coexist following a successful invasion under 39% of parameter combinations tested.

**Figure 2:**
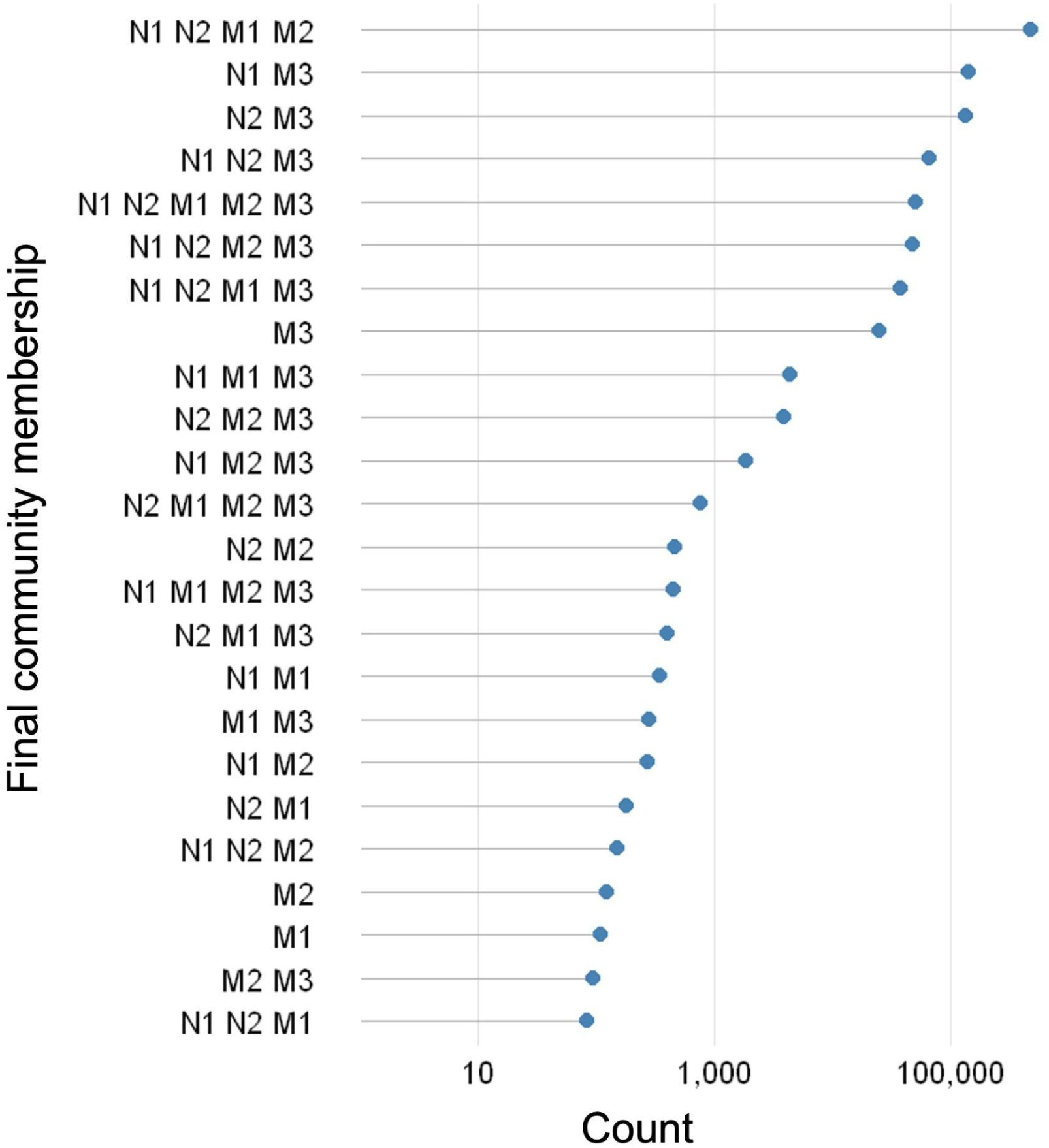
Counts of final community states across all simulations (note the log scale). State variables listed indicate the final community members, i.e. the entities that coexist following the introduction of *M*_3_ and return to equilibrium. For example, “N1 N2 M1 M2” indicates the invasion was resisted and the original community remains.

Fig. 3 demonstrates an example simulation in which only one host species and *M*_3_ remain. Prior to *M*_3_ introduction, host densities begin to stabilize by time 50 with *N*_1_ at a greater density than *N*_2_. Upon introduction of *M*_3_, the system’s microbiome composition is altered such that *M*_1_ is nearly extinct by time 90, at which point the density of *N*_1_ decreases, having lost the support of its associated microbiome (Fig. 3a and b). Ultimately this allows *N*_2_ to flourish and the system re-stabilizes with the opposite dominant host compared to pre-invasion, and *N*_1_ is driven extinct (Fig. 3c and d). Transitory dynamics following introduction of *M*_3_, such as those visible in Fig. 3a-b, are common and can even cause extinctions not only of resident species but also of *M*_3_ itself. These cases result in depauperate final communities containing neither *M*_3_ nor the full pre-invasion community (Fig. 2). While standard analytical methods cannot predict these scenarios, they represent only 0.17% of parameter combinations.

**Figure 3:**
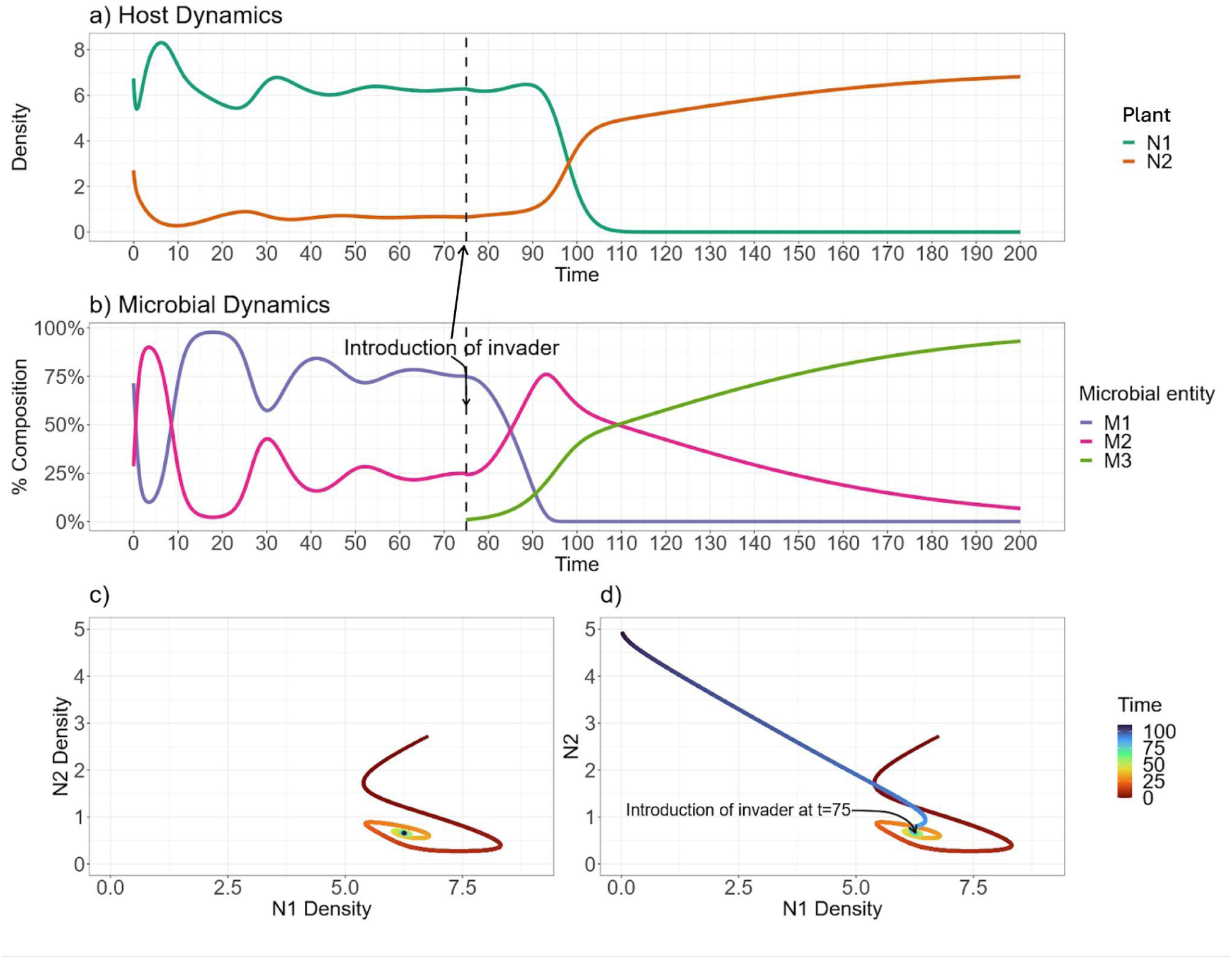
a) Example time series showing a change in host dynamics upon inoculation with a microbial invader *M*_3_ leading to extinction of host *N*_1_. b) Time series of the changing microbial community associated with the host dynamics presented in panel a. Pre-invasion, microbiome evenness shifts to stabilize host dynamics through negative feedback. Post-invasion, the microbial invader dominates and the stabilizing effect of the microbiome dynamics is compromised. c-d) Host densities over time for two systems with the same parameters. c) With no invasion, the community approaches a stable equilibrium in which both hosts are present, as described by the Bever et al. (1997) model. d) A microbe invades after 75 time-steps, sending the system to a new equilibrium in which *N*_1_ goes extinct.

The probability of successful *M*_3_ establishment is independent of pre-invasion microbiome or plant evenness (Fig. 4a and b). Likewise, the probability of any plant extinction following invasion is independent of initial microbiome or plant evenness (Fig. 4c and d). The data points at high probability values indicate the existence of parameter combinations that are particularly susceptible to plant extinction. Overall, invader persistence is more likely than plant extinction. However, the probability of successful establishment decreases when *M*_3_ invades later – specifically, as the timing of invasion approaches the time at which the community reaches its pre-invasion equilibrium, which is generally on the order of 100 time points (Fig. 5a, Fig. S2). Plants are also less likely to go extinct as the invader colonizes later, although the effect is weaker (Fig. 5b).

**Figure 4:**
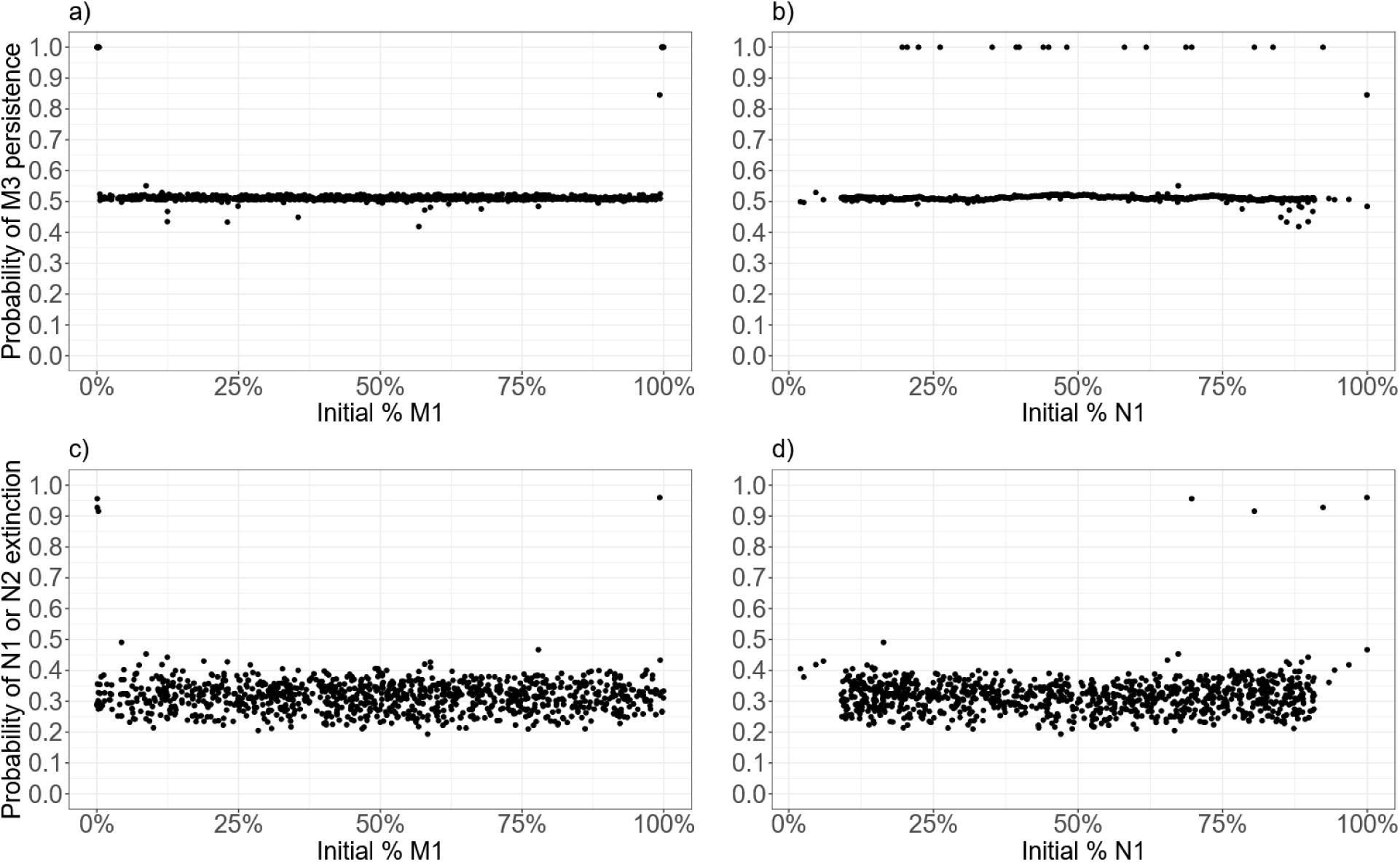
a-b) The probability of a successful invasion as a function of a) microbiome evenness pre-colonization and b) plant evenness pre-colonization. c-d) The probability of any plant extinction as a function of c) microbiome evenness pre-colonization and d) plant evenness pre-colonization. The x-axis represents evenness as the relative abundance of one plant or microbiome, because there are only two plant species and two resident microbiomes. For example, data points at M1=25% represent simulated invasions initiated with 25% *M*_1_ and 75% *M*_2_. Individual points represent a probability averaged over 1000 combinations of invader parameters while resident system parameters are held constant.

**Figure 5:**
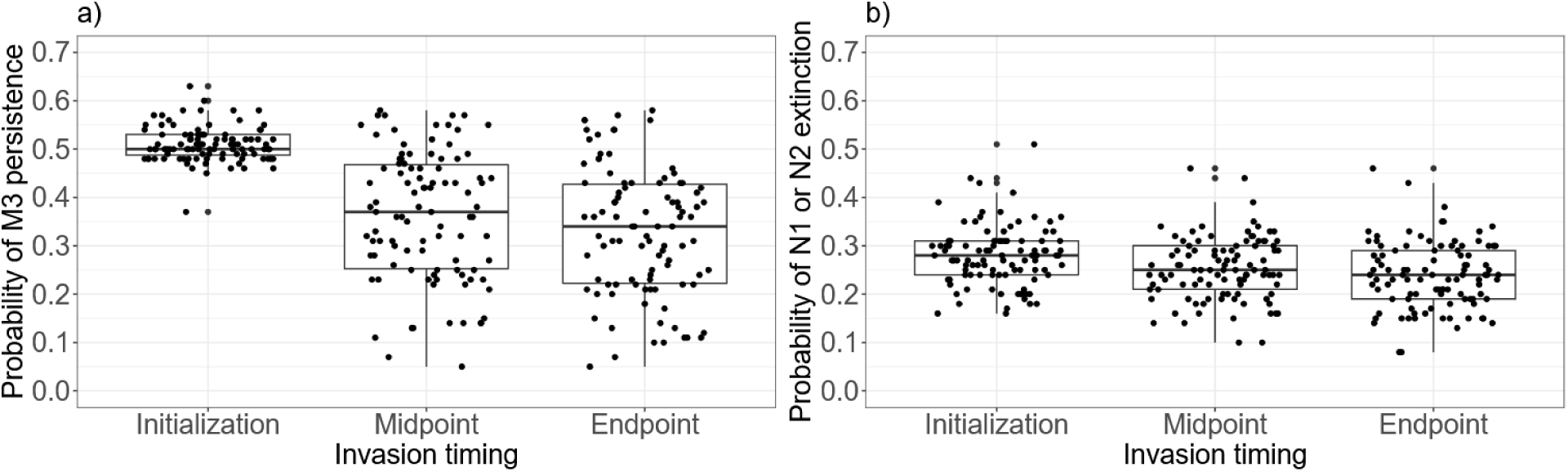
a) The probability of the persistence of the invading microbe *M*_3_ as a function of invasion timing. b) The probability of any host extinction as a function of invasion timing. Across both plots, individual points represent a probability averaged over 1000 combinations of invader parameters while resident system parameters are held constant.

The Routh-Hurwitz equilibrium stability analysis revealed three equilibrium categories following microbial colonization of the focal set of communities (i.e., the parameters chosen for equilibrium analysis). Either *M*_3_ is resisted and the system returns to its pre-invasion equilibrium, or *M*_3_ persists and either *N*_1_ or *N*_2_ determinatively goes extinct. These scenarios match the three most common scenarios found across our extensive parameter sampling conducted in the first phase of model analysis, suggesting that this targeted stability analysis is broadly representative (Fig. 2). Among parameter combinations resulting in successful invasions of the focal communities, the surviving plant species is determined by *α*_13_ and *α*_23_ (Fig. 6). In the case where *e*_1_ = *e*_2_ = 1, meaning both plants support *M*_3_ equally, *N*_2_ remains when *α*_13_ ≤ -1 and/or *α*_23_ < 0.75*α*_13_, meaning that either *M*_3_ has a significantly negative effect on *N*_1_ or that *M*_3_’s effect on *N*_2_ is more positive than its effect on *N*_1_ scaled by a factor of ∼0.75. Conversely, *N*_1_ instead remains if either *α*_13_ > -1, meaning *M*_3_ has a less negative effect on *N*_1_, or *α*_13_ < 1.75*α*_23_, meaning *M*_3_’s effect on *N*_1_ is more positive than its effect on *N*_2_ scaled by a factor of ∼1.75 (Fig. 6e). Interestingly, in this case there is also a sliver of parameter space between the above two outcomes where the invasion is resisted: when *α*_13_ > -1 and 1.33*α*_23_ ≲ *α*_13_ ≲ 1.75*α*_23_.

**Figure 6:**
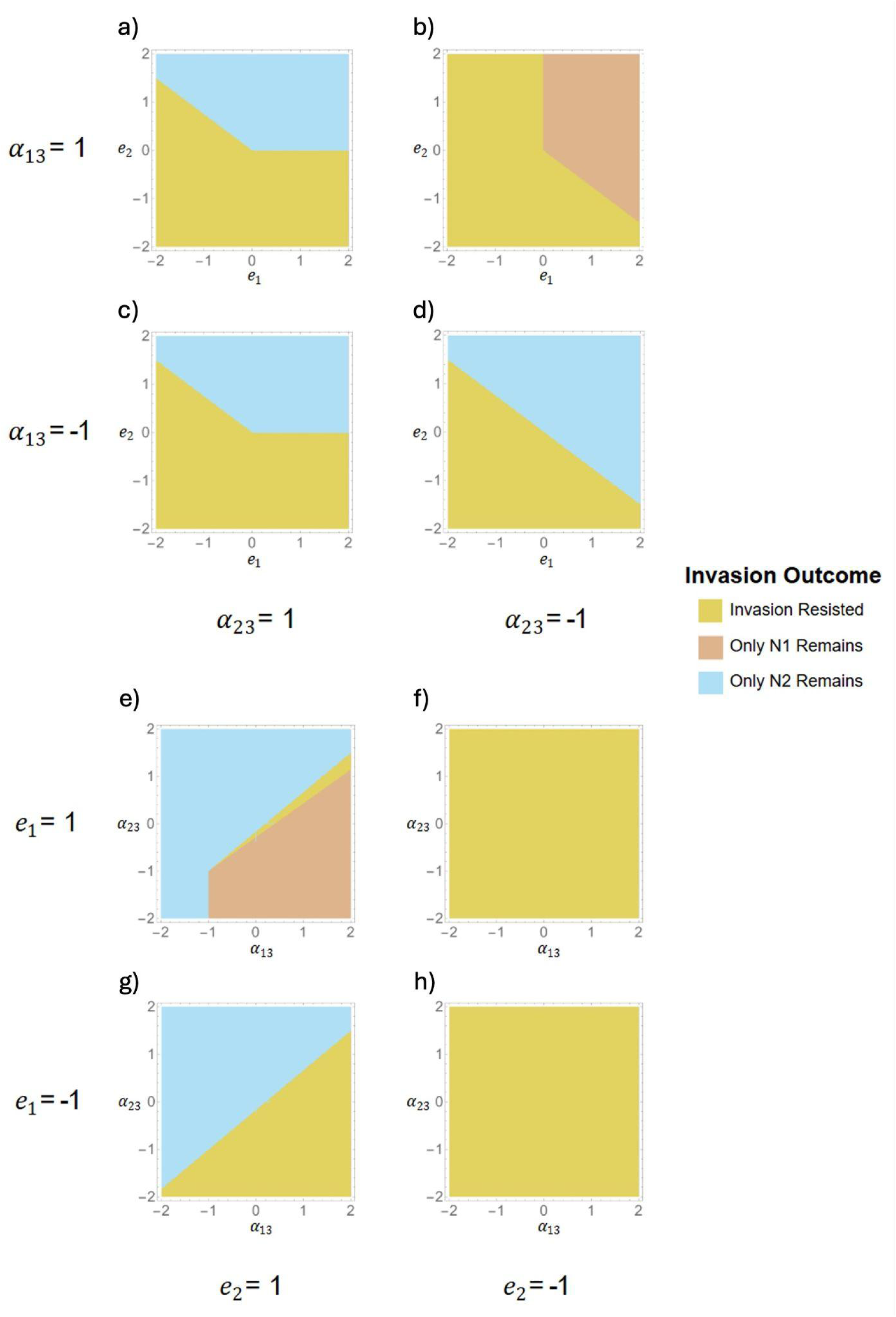
Phase diagrams relating invasion outcome to the plant-invader relationships. Results are displayed for a constant set of resident community parameters (*r*_1_=0.71, *r*_2_=1, *α*_11_=1, *α*_12_=1.8, *α*_21_=0.66, *α*_22_=1, *c*_11_=1, *c*_12_=0.86, *c*_21_=1, *c*_22_=1, and *v*=0.75). a-d) Panels displaying outcomes across all combinations of *e*_1_ and *e*_2_ (plant support of the invader) when the invader has a somewhat strong effect on plant growth rate, specifically *α*_13_=*α*_23_=|1|. e-h) Panels displaying outcomes across all combinations of *α*_13_ and *α*_23_ when the plants have a somewhat strong effect on the invader, specifically *e*_1_=*e*_2_=|1|.

## Discussion

We extend a version of the canonical plant-soil feedback model to examine the conditions allowing microbial invasions – including temporal dynamics – as well as the consequences of these invasions for plant-microbiome coexistence. Our model is sufficiently general to represent the invasion of any organism or group of organisms into a community of two plant species as long as the invader competes for resources with the resident microbiomes and interacts with the plants. For instance, the invader could be a generalist fungal pathogen or mutualist – or even a consortium of soil microbes – but our model is not intended to represent the transfer of a microbiome associated with a particular plant species. Our analyses suggest that invaders arriving during transient dynamics, i.e. while plant and microbiome densities are unstable, will be more likely to persist. Otherwise, our analyses suggest that plant-invader relationships, and not plant or microbiome evenness, determine invader success in our model. Given that an invader establishes in a community, it often interferes with the negative feedback between plant species and their microbiomes that enables plant coexistence, thereby causing local extinction of one of the plant species.

Our study demonstrates that the mechanism for plant coexistence in the classic plant-soil feedback model commonly breaks down when a new microbe successfully invades the system. This is not a trivial result given that our model includes intraspecific plant competition and therefore allows for robust coexistence. While the model originally proposed by Bever *et al*. (1997) predicts that two competing plants can coexist when microbiomes provide negative feedback on plant density, subsequent work pointed out that this model allows, at best, only tenuous plant coexistence which fails when additional plant species are added (Miller *et al*. 2022). Several approaches have been proposed to overcome this shortcoming and allow robust coexistence in multispecies plant-soil feedback communities (Abbott *et al*. 2021; Eppinga *et al*. 2018; Kulmatiski *et al*. 2011; Mack *et al*. 2019). We built our model as an extension of a model proposed by Abbott *et al*. (2021) that incorporates Lotka-Volterra competition terms including intraspecific competition, which can allow robust coexistence. Indeed, in our model the plants continue to coexist following successful invasions across 39% of the simulated communities. Therefore, it is also common that microbial invaders fail to disrupt plant coexistence in our model. Still, in the other 61% of simulated communities the invader interferes with the negative feedback among the plant species and their microbiomes and causes a plant extirpation. Given that microbial invasions are not likely to be rare, additional mechanisms may be needed to account for plant coexistence in nature such as trait trade-offs, microbially-mediated fitness differences, or metacommunity processes (Castledine *et al*. 2020; Kandlikar *et al*. 2019; Thompson *et al*. 2020).

Neither plant extinction nor invader success depend on initial microbiome or plant evenness in our model, but invader success is highest during transient dynamics. The greater importance of biotic interactions compared to resident evenness in determining microbial invasions is consistent with theory. While more functionally diverse microbial communities better resist microbial colonization in environments with many niches, colonization under low niche diversity – akin to our model, which did not include niche diversity – is determined by the resident community composition (Eisenhauer *et al*. 2016). Additionally, our finding that colonization was most likely during transient dynamics is consistent with dynamical systems theory (Strogatz 2015). As a greater amount of time passes before the introduction of the invader, the resident system travels farther into the basin of attraction of its equilibrium, meaning the perturbation caused by the invader must be larger to send the system to an alternate equilibrium incorporating the invader. Therefore, community assembly patterns are different during transient states, including higher rates of establishment of new species and higher beta diversity, compared to patterns under stable states (Dooley *et al*. 2024; Fukami & Nakajima 2011). It follows, then, that higher rates or intensities of disturbance – which perturb communities from their equilibrium and lead to transient dynamics – will facilitate microbial invasions. Indeed, experimental evidence shows that environmental variability can facilitate invasion (Tucker & Fukami 2014). A major challenge moving forward will be mapping transient dynamics theory to empirical systems, especially given that transient states last long compared to immigration rates and community dynamics (Fukami & Nakajima 2011).

Existing studies on the temporal dimension of plant-soil microbiome interactions have primarily focused on the demographic processes involved in soil microbial conditioning and plant response phases. For example, patch-occupancy models have been used to study the effects of soil microbial conditioning on plant establishment (Ke & Levine 2021; Krishnadas & Stump 2021; Miller & Allesina 2021). Theoreticians have also modeled microbial effects on plant demographic processes. For example, Mordecai (2013) found that a fungal pathogen shared by an invader annual and native perennial plant differently impacted seed infection rates, germination rates, and adult host survival. Contrary to expectation, pathogen spillover in multi-host communities had a broad range of outcomes including coexistence, exclusion, and priority effects (Mordecai 2013a). Subsequent work in the same study system found that foliar fungal pathogens did not promote negative frequency dependence and plant coexistence (Spear & Mordecai 2018). However, most research on community assembly in the context of plant communities focuses on the plant species, and few studies have investigated the community-level consequences of microbial invasions and their timing (Dooley *et al*. 2024; Peay 2018; Rivett *et al*. 2018).

Our stability analysis indicates that beyond invasion timing, the plant-invader relationships determine invader success. Since the differences between *N*_1_ and *N*_2_ are arbitrary and entirely due to chosen parameters, the apparent difference in importance of *e*_1_ and *e*_2_ on invader success in this set of communities is due to the particular parameters used in this stability analysis. For example, *N*_2_ has a higher intrinsic rate of increase, which could enable it to reach higher densities, meaning that when its effect on *M*_3_ is negative (*e*_2_ < 0) it can more effectively prevent *M*_3_ from taking hold. Here we should note the limitations of the stability analysis: namely, we varied *e*_1_, *e*_2_, *α*_13_, and *α*_23_ for fixed values of all other system parameters. Repeating the analysis while allowing one other parameter to vary yielded results in agreement with those presented. However, it was not feasible for us to conduct the same analyses with all system parameters left free.

There are many important sources of variation in real ecosystems which we were not able to explicitly investigate within the scope of our study. One such source of variation is microbiome type. Theory suggests that microbiomes with a more facilitative effect on their host plant – e.g. dominated by mycorrhizae – encourage priority effects and lower plant diversity, while microbiomes with a more negative effect – e.g. dominated by pathogens – encourage plant coexistence (Bever *et al*. 1997; Crawford *et al*. 2019; Umbanhowar & McCann 2005). While we did not assume neutral effects of microbiomes on their hosts, we randomized these effects and did not focus our study on the influence of their sign. Therefore, further research could more explicitly investigate different patterns expected when positive or negative microbiome effects dominate. Different microbial functional groups such as mycorrhizae or pathogens can also exhibit different community assembly patterns. For example, some functional groups can be more strongly structured by dispersal while others are more strongly structured by environmental filtering, although studies disagree on the particular hierarchy (Gacura *et al*. 2024; Liang *et al*. 2023; Wang *et al*. 2022). Assembly patterns are also likely to depend on spatial scale, where priority effects are more likely to be important at small scales and factors such as dispersal can be more important at larger scales. For example, using an approach focusing on spatial effects, Gacura *et al*. (2024) were able to explain the assembly of fungal functional groups across forest stands but not at small scales, citing priority effects as a likely explanation. Our results may therefore be most applicable at small spatial scales. Further research is needed to extend our approach to incorporate metacommunity processes and community coalescence (Castledine *et al*. 2020; Thompson *et al*. 2020).

Our study contributes to a new node of ecological theory at the intersection of multispecies coexistence, host-microbiome dynamics, and time-dependent interactions, all of which appear to be important for understanding and predicting ecosystem dynamics (Zou *et al*. 2024). While many previous studies have either assumed all species are initially present or assume an invader enters a system at equilibrium, in reality communities rarely appear to be in a demonstrable equilibrium. Plant communities, including their microbiomes, are frequently undergoing long-term succession as well as phenological changes. Meanwhile, individual plants and microbes are constantly undergoing development and senescence. Therefore, microbial invasions may well have differential impacts on plant communities depending on their timing with respect to plant succession and development. It is increasingly important to understand how species interactions change over successional and phenological timelines as ecosystems are recovering more frequently from anthropogenic disturbances and interventions, and as climate change creates phenological mismatches among interacting species. In the years to come, research in this emerging field of complex time-dependent interactions among multiple species and microbiomes has the potential to provide managers with the knowledge necessary to become stewards of biodiversity and ecosystem services in a new era of species migrations and global change.

## Acknowledgements

The authors would like to thank Corbin Jones and members of the Yitbarek Lab (UNC), Gibert Lab (Duke University), Leibold Lab (University of Florida), and the Theoretical Ecology and Evolution Group at UNC for feedback on the manuscript. The authors acknowledge seminar invitations from North Carolina State University and Yale University where ideas for this manuscript were presented and feedback was received. The authors acknowledge funding from the UNC Summer Undergraduate Research Fellowship (WHP) and UNC Junior Faculty Award (SY).

## Author contributions

WHP and SY planned and designed the research. WHP analyzed the model, and WHP and CJR produced the figures. WHP, SY, and CJR wrote the manuscript.

## Data accessibility

The code used in this study is openly available in Zenodo at <https://doi.org/10.5281/zenodo.17251424>.

## Supplemental Information

**Figure S1:**
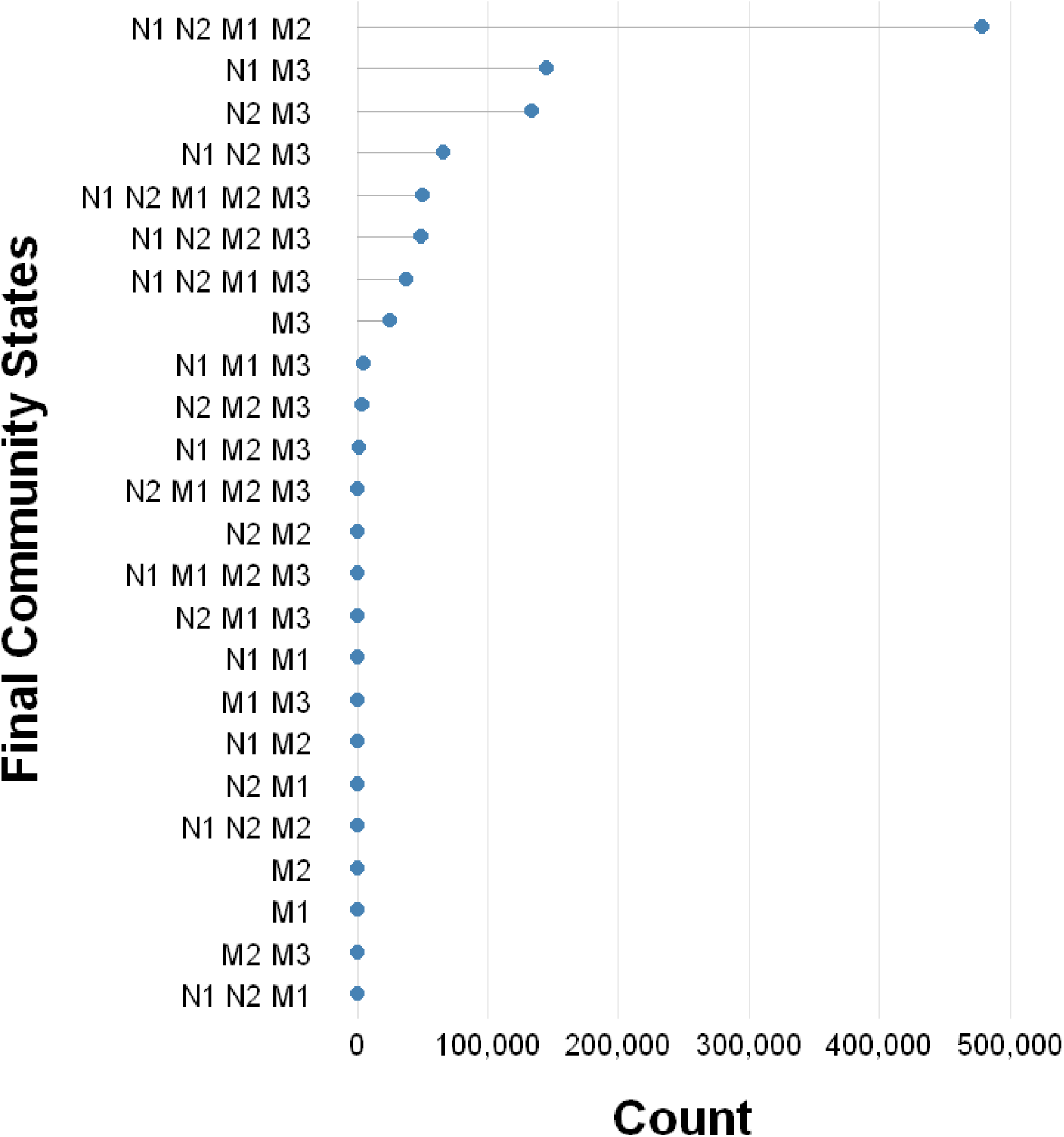
Counts of final community states across all simulations. State variables listed indicate the final community members, i.e. the entities that coexist following the introduction of *M*_3_ and return to equilibrium. For example, “N1 N2 M1 M2” indicates the invasion was resisted and the original community remains.

**Figure S2:**
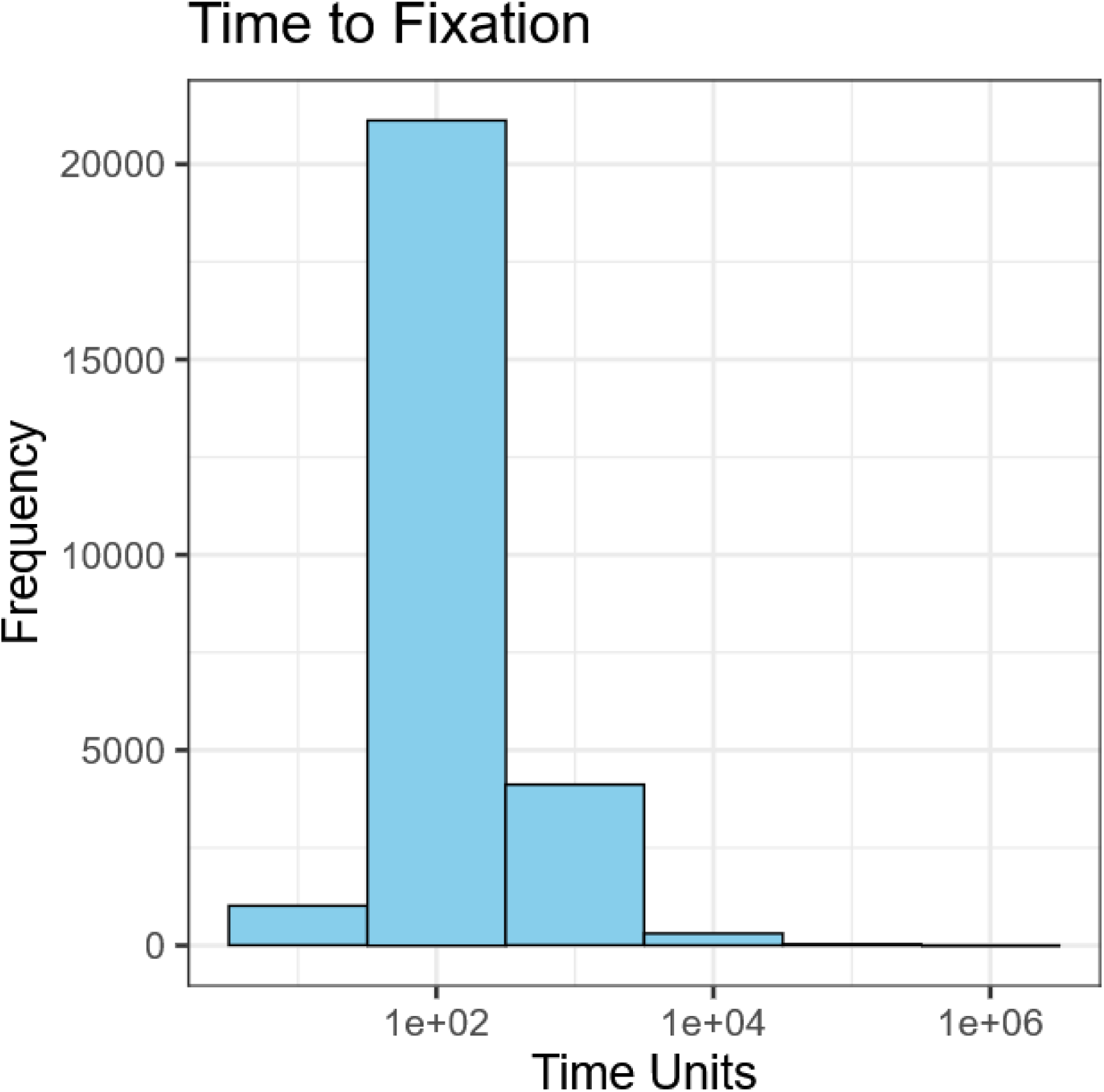
Histogram of time steps required to reach equilibrium for various combinations of resident community parameters sampled in our simulations. The time to fixation depends on the system parameters, but a fixation time on the order of 100 time units was most frequently observed.

## References

Abbott, K.C., Eppinga, M.B., Umbanhowar, J., Baudena, M. & Bever, J.D. (2021). Microbiome influence on host community dynamics: Conceptual integration of microbiome feedback with classical host–microbe theory. Ecol. Lett., 24, 2796–2811.

Adler, P.B., HilleRisLambers, J. & Levine, J.M. (2007). A niche for neutrality. Ecol. Lett., 10, 95–104.

Albright, M.B.N., Sevanto, S., Gallegos-Graves, L.V. & Dunbar, J. (2020). Biotic Interactions Are More Important than Propagule Pressure in Microbial Community Invasions. mBio, 11, e02089–20.

Bever, J.D. (2003). Soil community feedback and the coexistence of competitors: conceptual frameworks and empirical tests. New Phytol., 157, 465–473.

Bever, J.D., Mangan, S.A. & Alexander, H.M. (2015). Maintenance of Plant Species Diversity by Pathogens. Annu. Rev. Ecol. Evol. Syst., 46, 305–325.

Bever, J.D., Westover, K.M. & Antonovics, J. (1997). Incorporating the Soil Community into Plant Population Dynamics: The Utility of the Feedback Approach. J. Ecol., 85, 561–573.

Carnell, R. (2022). lhs: Latin Hypercube Samples.

Castledine, M., Sierocinski, P., Padfield, D. & Buckling, A. (2020). Community coalescence: an eco-evolutionary perspective. Philos. Trans. R. Soc. B Biol. Sci., 375, 20190252.

Chaudhary, V.B., Aguilar-Trigueros, C.A., Mansour, I. & Rillig, M.C. (2022). Fungal Dispersal Across Spatial Scales. Annu. Rev. Ecol. Evol. Syst., 53, 69–85.

Crawford, K.M., Bauer, J.T., Comita, L.S., Eppinga, M.B., Johnson, D.J., Mangan, S.A., et al. (2019). When and where plant-soil feedback may promote plant coexistence: a meta-analysis. Ecol. Lett., 22, 1274–1284.

Debray, R., Herbert, R.A., Jaffe, A.L., Crits-Christoph, A., Power, M.E. & Koskella, B. (2022). Priority effects in microbiome assembly. Nat. Rev. Microbiol., 20, 109–121.

Dooley, K.D., Henry, L.P. & Bergelson, J. (2024). Impact of timing on the invasion of synthetic bacterial communities. ISME J., 18, wrae220.

Eisenhauer, N., Barnes, A.D., Cesarz, S., Craven, D., Ferlian, O., Gottschall, F., et al. (2016). Biodiversity-ecosystem function experiments reveal the mechanisms underlying the consequences of biodiversity change in real world ecosystems. J. Veg. Sci., 27, 1061–1070.

van Elsas, J.D., Chiurazzi, M., Mallon, C.A., Elhottovā, D., Krištůfek, V. & Salles, J.F. (2012). Microbial diversity determines the invasion of soil by a bacterial pathogen. Proc. Natl. Acad. Sci., 109, 1159–1164.

Elton, C.S. (2020). The Ecology of Invasions by Animals and Plants. Springer International Publishing, Cham.

Eppinga, M.B., Baudena, M., Johnson, D.J., Jiang, J., Mack, K.M.L., Strand, A.E., et al. (2018). Frequency-dependent feedback constrains plant community coexistence. Nat. Ecol. Evol., 2, 1403–1407.

Finlay, B.J. & Clarke, K.J. (1999). Ubiquitous dispersal of microbial species. Nature, 400, 828–828.

Fukami, T. & Nakajima, M. (2011). Community assembly: alternative stable states or alternative transient states?: Alternative transient states. Ecol. Lett., 14, 973–984.

Gacura, M.D., Zak, D.R. & Blackwood, C.B. (2024). From individual leaves to forest stands: importance of niche, distance decay, and stochasticity vary by ecosystem type and functional group for fungal community composition. FEMS Microbiol. Ecol., 100, fiae016.

Hannula, S.E., Ma, H., Pérez-Jaramillo, J.E., Pineda, A. & Bezemer, T.M. (2020). Structure and ecological function of the soil microbiome affecting plant–soil feedbacks in the presence of a soil-borne pathogen. Environ. Microbiol., 22, 660–676.

Hindmarsh, A.C. & Petzold, L.R. (2005). LSODA, Ordinary Differential Equation Solver for Stiff or Non-Stiff System.

Holt, R. & Lawton, J. (2003). The Ecological Consequences of Shared Natural Enemies. Annu. Rev. Ecol. Syst., 25, 495–520.

Jeschke, J.M., Bacher, S., Blackburn, T.M., Dick, J.T.A., Essl, F., Evans, T., et al. (2014). Defining the Impact of Non-Native Species. Conserv. Biol., 28, 1188–1194.

Jiang, J., Abbott, K.C., Baudena, M., Eppinga, M.B., Umbanhowar, J.A. & Bever, J.D. (2020). Pathogens and Mutualists as Joint Drivers of Host Species Coexistence and Turnover: Implications for Plant Competition and Succession. Am. Nat., 195, 591–602.

Kandlikar, G.S., Johnson, C.A., Yan, X., Kraft, N.J.B. & Levine, J.M. (2019). Winning and losing with microbes: how microbially mediated fitness differences influence plant diversity. Ecol. Lett., 22, 1178–1191.

Ke, P.-J. & Levine, J.M. (2021). The Temporal Dimension of Plant-Soil Microbe Interactions: Mechanisms Promoting Feedback between Generations. Am. Nat., 198, E80–E94.

Krishnadas, M. & Stump, S.M. (2021). Dispersal limitation and weaker stabilizing mechanisms mediate loss of diversity with edge effects in forest fragments. J. Ecol., 109, 2137–2151.

Kulmatiski, A., Heavilin, J. & Beard, K.H. (2011). Testing predictions of a three-species plant–soil feedback model. J. Ecol., 99, 542–550.

Liang, J., Ding, Z., Lie, G., Zhou, Z., Zhang, Z. & Hu, H. (2023). Patterns and drivers of phylogenetic diversity of seed plants along an elevational gradient in the central Himalayas. Glob. Ecol. Conserv., 47, e02661.

Litvak, Y. & Bäumler, A.J. (2019). The founder hypothesis: A basis for microbiota resistance, diversity in taxa carriage, and colonization resistance against pathogens. PLoS Pathog., 15, e1007563.

Mack, K.M.L., Eppinga, M.B. & Bever, J.D. (2019). Plant-soil feedbacks promote coexistence and resilience in multi-species communities. PLOS ONE, 14, e0211572.

Madi, N., Vos, M., Murall, C.L., Legendre, P. & Shapiro, B.J. (2020). Does diversity beget diversity in microbiomes? eLife, 9, e58999.

Mangan, S.A., Schnitzer, S.A., Herre, E.A., Mack, K.M.L., Valencia, M.C., Sanchez, E.I., et al. (2010). Negative plant–soil feedback predicts tree-species relative abundance in a tropical forest. Nature, 466, 752–755.

Miller, Z.R. & Allesina, S. (2021). Metapopulations with habitat modification. Proc. Natl. Acad. Sci. U. S. A., 118, e2109896118.

Miller, Z.R., Lechón-Alonso, P. & Allesina, S. (2022). No robust multispecies coexistence in a canonical model of plant–soil feedbacks. Ecol. Lett., 25, 1690–1698.

Mordecai, E.A. (2013a). Consequences of Pathogen Spillover for Cheatgrass-Invaded Grasslands: Coexistence, Competitive Exclusion, or Priority Effects. Am. Nat., 181, 737–747.

Mordecai, E.A. (2013b). Despite spillover, a shared pathogen promotes native plant persistence in a cheatgrass-invaded grassland. Ecology, 94, 2744–2753.

Parker, I.M. & Gilbert, G.S. (2004). The Evolutionary Ecology of Novel Plant-Pathogen Interactions. Annu. Rev. Ecol. Evol. Syst., 35, 675–700.

Peay, K.G. (2018). Timing of mutualist arrival has a greater effect on *Pinus muricata* seedling growth than interspecific competition. J. Ecol., 106, 514–523.

Pedersen, T.L. (2025). patchwork: The Composer of Plots.

R Core Team. (2022). R: A language and environment for statistical computing.

Rivett, D.W., Jones, M.L., Ramoneda, J., Mombrikotb, S.B., Ransome, E. & Bell, T. (2018). Elevated success of multispecies bacterial invasions impacts community composition during ecological succession. Ecol. Lett., 21, 516–524.

Senthilnathan, A. & D’Andrea, R. (2024). Coexistence of Competing Plants Under Plant–Soil Feedback. Ecol. Lett., 27, e14503.

Slowikowski, K. (2024). ggrepel: Automatically Position Non-Overlapping Text Labels with “ggplot2.”

Soetaert, K., Petzoldt, T. & Setzer, R.W. (2010). Solving Differential Equations in R: Package deSolve. J. Stat. Softw., 33, 1–25.

Spear, E.R., Coley, P.D. & Kursar, T.A. (2015). Do pathogens limit the distributions of tropical trees across a rainfall gradient? J. Ecol., 103, 165–174.

Spear, E.R. & Mordecai, E.A. (2018). Foliar pathogens are unlikely to stabilize coexistence of competing species in a California grassland. Ecology, 99, 2250–2259.

Strogatz, S.H. (2015). Nonlinear dynamics and chaos: with applications to physics, biology, chemistry, and engineering. 2nd Edition. Chapman and Hall/CRC, New York.

Thompson, P.L., Guzman, L.M., De Meester, L., Horváth, Z., Ptacnik, R., Vanschoenwinkel, B., et al. (2020). A process-based metacommunity framework linking local and regional scale community ecology. Ecol. Lett., 23, 1314–1329.

Tucker, C.M. & Fukami, T. (2014). Environmental variability counteracts priority effects to facilitate species coexistence: evidence from nectar microbes. Proc. R. Soc. B Biol. Sci., 281, 20132637.

Umbanhowar, J. & McCann, K. (2005). Simple rules for the coexistence and competitive dominance of plants mediated by mycorrhizal fungi. Ecol. Lett., 8, 247–252.

Wang, C., Ma, L., Zuo, X., Ye, X., Wang, R., Huang, Z., et al. (2022). Plant diversity has stronger linkage with soil fungal diversity than with bacterial diversity across grasslands of northern China. Glob. Ecol. Biogeogr., 31, 886–900.

Wickham, H., Averick, M., Bryan, J., Chang, W., McGowan, L.D., François, R., et al. (2019). Welcome to the Tidyverse. J. Open Source Softw., 4, 1686.

Wolfram Research, Inc. (2022). Mathematica.

Zou, H.-X., Yan, X. & Rudolf, V.H.W. (2024). Time-dependent interaction modification generated from plant–soil feedback. Ecol. Lett., 27, e14432.

